# Does white matter structure relate to hemispheric language lateralisation? A systematic review

**DOI:** 10.1101/2025.01.04.631320

**Authors:** Ieva Andrulyte, Eszter Demirkan, Francesca M. Branzi, Laura J. Bonnett, Simon S. Keller

## Abstract

The relationship between functional language lateralisation and diffusion MRI-based white matter metrics remains a subject of considerable interest and complexity. This systematic review aims to synthesise existing diffusion MRI studies examining white matter correlates of functional language dominance. Twenty-five studies were identified through searches of Web of Science, Scopus, and Ovid MEDLINE (search period: inception to 16th March 2023) involving adults with epilepsy, tumours, or healthy controls. The results suggest that while the arcuate fasciculus, particularly its fractional anisotropy (FA), is commonly associated with language lateralisation in clinical populations, the findings in healthy individuals are more variable, often influenced by factors such as handedness. Other white matter tracts, including the corpus callosum and uncinate fasciculus, showed less consistent associations with language dominance across studies. Interestingly, temporal lobe regions, especially those involved in semantic processing, exhibited stronger correlations with diffusion measures compared to areas associated with phonological tasks. Methodological inconsistencies, such as variability in sample selection, task design, and analytical techniques, were identified as significant challenges in comparing findings across studies. Future research should aim for larger, more diverse sample sizes, whole-brain approaches, and a wider range of fMRI tasks to better elucidate the role of white matter in language lateralisation. If regions of interest (ROI)-based studies are utilised, a more standardised approach to tract segmentation should be adopted to ensure consistency and improve comparability across studies.

## Introduction

Hemispheric structural asymmetry, a fundamental anatomic characteristic of the human brain, refers to the differences in structure between the left and right hemispheres (Gazzaniga, 2000; Kong et al., 2020). Despite decades of research, the mechanisms underlying this asymmetry are not fully understood (Badzakova-Trajkov et al., 2016).

Functional hemispheric lateralisation, particularly for language, was first recognised in the 19th century and has since been extensively studied through functional magnetic resonance imaging (fMRI) (Esteves et al., 2020). In around 95% of right-handed individuals, language functions are predominantly lateralised to the left hemisphere, while the remainder show right-hemisphere or bilateral representation (Corballis et al., 2012). The mechanisms behind this distribution are still unclear, though it is hypothesised that brain structure might support functional lateralisation (Keller et al., 2018).

The anatomical basis for language lateralisation has traditionally been linked to structural differences in specific brain regions, such as the planum temporale (Josse et al., 2003) and inferior frontal gyrus (Josse et al., 2009). However, advances in neuroimaging, particularly diffusion tensor imaging (DTI), have shifted attention towards the role of white matter connectivity in supporting language function. DTI enables researchers to explore various aspects of white matter tracts, such as fractional anisotropy (FA), volume, and streamline count, offering insights into how these pathways support language processing and lateralisation (Astrakas, 2023). Recent studies have emphasised the importance of specific white matter tracts, including the arcuate fasciculus, uncinate fasciculus, and frontal aslant tract, in supporting language functions, with damage to these tracts being linked to language deficits, particularly aphasia (Basilakos et al., 2014; Geller et al., 2019; Hope et al., 2016; Kim et al., 2019; Marchina et al., 2011; Wang et al., 2013).

Despite the wealth of literature on hemispheric asymmetry and language processing, systematic reviews addressing the relationship between structure and function are scarce. Existing reviews primarily focus on structural asymmetry (Esteves et al., 2020; Güntürkün et al., 2020; Hervé et al., 2013), its relation to cognition (Hartwigsen et al., 2021), and applications in surgical assessments (Duncan et al., 2016), or examine language-related white matter tracts in isolation (Friederici, 2009). Systematic reviews of language lateralisation have primarily concentrated on fMRI studies (Bauer et al., 2014; Bradshaw, Bishop, et al., 2017; Bradshaw, Thompson, et al., 2017), typically without consideration of the relationships between structural and functional imaging modalities. The challenge of comparing findings across different fMRI studies alone is compounded by variations in methodologies, including differences in participant demographics, language tasks, and imaging techniques employed (Costafreda et al., 2006). Furthermore, the integration of diffusion MRI with functional assessments poses additional methodological challenges, as each imaging modality has distinct protocols and definitions of anatomical regions.

To address this gap, we performed a systematic review to evaluate the relationship between white matter diffusion metrics (i.e. FA, volume, number of streamlines) and language lateralisation across diverse populations and methodologies. By examining the correlation between diffusion metrics and language dominance, we seek to determine how variations in white matter architecture may relate to language lateralisation across different populations, including clinical and non-clinical groups.

## Methods

To address the objectives of this research, a systematic review was conducted following the Preferred Reporting Items for Systematic Reviews and Meta Analyses (PRISMA) statement (Page et al., 2021) (Supplementary Table 1). This review was preregistered with PROSPERO on May 3rd, 2023 (Registration number CRD42023402015). The team of reviewers comprised five individuals (I.A. = I. Andrulyte, E.D. = E. Demirkan, F.M.B. = F. M. Branzi, L.B. = L. Bonnett, S.K. = S. Keller).

### Search strategy

We conducted a systematic search to investigate the relationship between diffusion-based MRI measures of white matter and functional language dominance, specifically examining differences in among individuals with left-hemispheric, right-hemispheric, and bihemispheric language representation. The search commenced on 17 April 2023 and was finalised on 16 May 2023. To identify relevant studies, we used three online databases: MEDLINE, Scopus, and Web of Science. We focused on studies with full text availability, published in English, without imposing any year restrictions.

Our search strategy employed multiple concepts with alternative terms spanning various related domains. Concept 1 concentrated on language lateralization in the brain, using search terms such as “language lateral*”, “language dominance,” and “language asymmetr*.” Concept 2 addressed hemispheric function, incorporating terms like “hemispheric dominance,” “hemispheric asymmetr*,” and “brain asymmetr*.” These two concepts were combined using Boolean operators to identify studies focused on language function within the framework of brain lateralization (e.g., combining “language” with terms related to hemispheric functioning). Concept 3 focused on brain imaging methodologies, particularly Diffusion Tensor Imaging (DTI) and related techniques (search terms: “DTI,” “diffusion tensor imaging,” “tractography,” “white matter,” “fibre,” “fiber”). This step integrated terms related to brain lateralization and hemispheric dominance with advanced imaging techniques to locate relevant studies. The final step involved integrating these concepts into a comprehensive search strategy (Supplementary Table 2).

All references identified through this search were imported into the Covidence system by I.A., where duplicates were removed through a combination of software and manual review. After eliminating duplicates, the search resulted in a total of 687 references. A backup of the initial search was saved in Excel for reference. All reviewers had access to the same article list; however, the review process was blinded. Title and abstract screening were performed in Covidence, with selected articles exported to Excel for full- text screening

### Inclusion/Exclusion criteria

Two reviewers (I.A. and E.D.) independently assessed the title and abstracts of 687 records to select studies that met the criteria. In cases of disagreement, a third reviewer (F.M.B.) conducted an independent evaluation. The following eligibility criteria were applied:

1. Study type: We included all studies except letters, abstracts, case series, and reviews.
2. Participants: Studies involving adults (as defined by the study) with epilepsy, tumours, or healthy controls were included.
3. MRI modalities: Only imaging studies incorporating both diffusion MRI and (1) fMRI, (2) WADA, or (3) dichotic listening measurements were considered.
4. Functional laterality: Only studies showing the distribution of functional laterality were considered.
5. Data analysis: Studies examining structure-function relationships between language lateralisation and structural correlates were included.
6. Availability: Studies without full-text access were excluded.
7. Language: Studies not written in English were excluded.

Following this screening process, we excluded all records that did not meet the above criteria (n=632). Articles that were included based on the title and abstract screening (n=54) were retrieved for independent screening by I.A. and E.D. in Excel (Supplementary Table 3). Out of the 54 studies, 25 met the eligibility criteria and were included in our review. Two studies were excluded at this stage: one due to a language barrier (written in German; Rothenberger & Roessner, 2006), and another because it was a review article (Ocklenburg et al., 2016). The remaining 28 studies were excluded as they either lacked sufficient detail or were considered outside the scope of this review. A flow diagram outlining the study selection process is provided in Figure 1.

**Figure 1.**
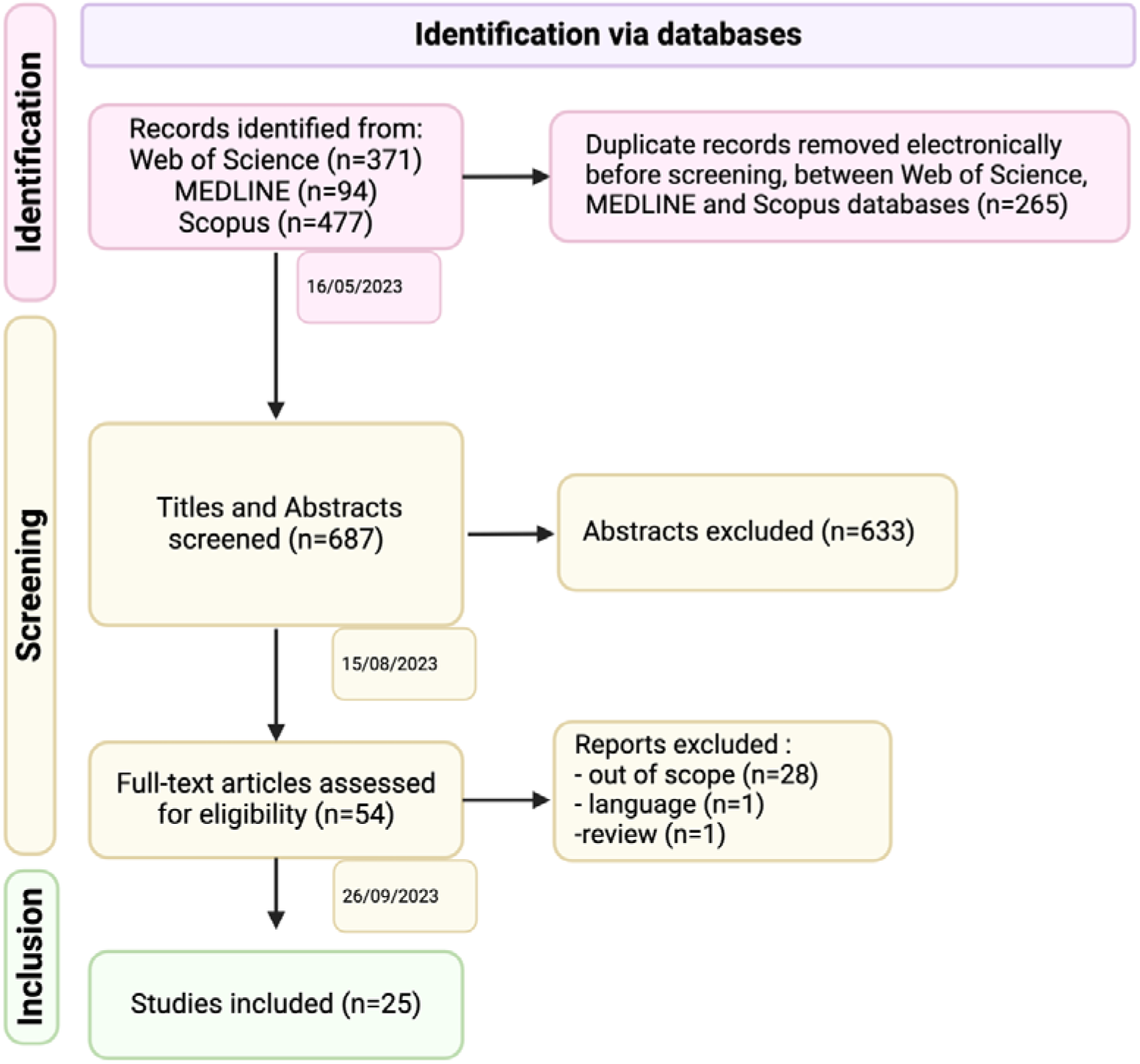
Systematic review flow diagram (PRISMA 2020 template).

### Data extraction and analysis

From the 25 studies included in our review, we extracted the following information:

1. Participant details: Information on participants involved in the study, including sample size, gender ratio, average age, health status, handedness, and native language.
2. Methodological details: Information on MRI strength (Tesla), the method used to assess functional language dominance (fMRI, WADA, dichotic listening), language tasks, baseline conditions for language tasks (if fMRI was used), regions-of-interest (ROIs) used to determine language dominance, thresholds for categorising laterality groups, categories used for language dominance (e.g., typical/atypical, lateralised/bilateral, left/right/bilateral), diffusion MRI methods (software, probabilistic/deterministic tools), diffusion-based MRI measures examined (e.g., volume), the approach used for diffusion MRI (i.e., ROI or whole brain), specific tracts studied (if ROI-based), statistical tests used to examine relationships, and covariates in the model.
3. Results: Information on language dominance (number of individuals in each laterality group) and findings regarding the relationships between language lateralisation and white matter tracts.

Two reviewers (I.A. and E.D.) were responsible for extracting this information. I.A. extracted data from all studies, and E.D. independently extracted data from 16% of the studies (4 studies) to check for agreement. There were no discrepancies between the reviewers. The aforementioned information is available in Supplementary Table 4.

For data extraction and analysis, we first grouped the studies by population (healthy vs. diseased). Next, studies were categorised based on white matter tracts, including the arcuate fasciculus, superior longitudinal fasciculus, inferior longitudinal fasciculus, corpus callosum, corticospinal fibres, corticobulbar tract, uncinate fasciculus, inferior fronto-occipital fasciculus, frontal aslant tract, and optic radiation. For synthesis and presentation, experiments were further grouped by diffusion metrics—microstructural measures (e.g., FA, mean diffusivity (MD)), which capture properties like water diffusion within voxel, and macrostructural measures (e.g., tract volume, fibre density), which assess the overall structure of white matter pathways rather than their individual components. Studies were also grouped by language tasks (phonology and semantics) to reduce heterogeneity in the reported results. Language tasks were further classified by difficulty, considering whether the task was passive, required a yes/no response, or was multiple choice or generative, and compared against the baseline type (active or passive). The results of all studies were compiled in an Excel file, and we synthesised group comparisons (left vs. right vs. bilateral) and differences between conditions in relation to the results.

We used narrative synthesis to summarise the evidence from the included studies, described the findings in the upcoming sections, and presented them in figures and tables. Due to substantial variability in language tasks, ROIs used to determine laterality, diffusion metrics, and tractography tools, performing a meta-analysis was not feasible.

### Risk of bias assessment criteria

We used the modified Newcastle-Ottawa Scale (NOS) to evaluate the methodological quality of cross-sectional studies (Herzog et al., 2013) (Supplementary Table 5). The NOS assesses three distinct domains: selection, comparability, and outcome; each of them having a question in different subsections. We removed the questions associated with treatment outcomes because it was not a question of interest in our study and, based on the latest findings in laterality research, extended this tool by incorporating features about missing data, lesion location, statistical methods, and data interpretation (Bidula et al., 2017; Pasquini et al., 2022; Vingerhoets et al., 2023).

The scale is graded out of 11 points, with 10–11 points indicating a study of good quality, 6–9 points indicating a fair-quality study, and 0–5 points indicating a poor-quality study (Wong et al., 2023). Quality appraisal was conducted independently by two reviewers (I.A. and E.D.). Their scores were compared, and any discrepancies were resolved by consensus.

## Results (study descriptions)

### Included studies

A total of 687 studies were initially identified. After excluding duplicates and studies that were out of scope, published in languages different from English, as well as reviews, clinical cases, letters, studies involving non-adult participants (children or adolescents), and further irrelevant studies, 25 studies were retained for final analysis (Figure 1). The publication dates of these included studies ranged from November 2003 to December 2022 (Figure 2). All studies were cross-sectional in design, with sample sizes varying between 9 and 89 participants. Fifteen studies employed 3-Tesla scanners, while the remaining 10 used 1.5-Tesla scanners (Supplementary Table 4).

**Figure 2.**
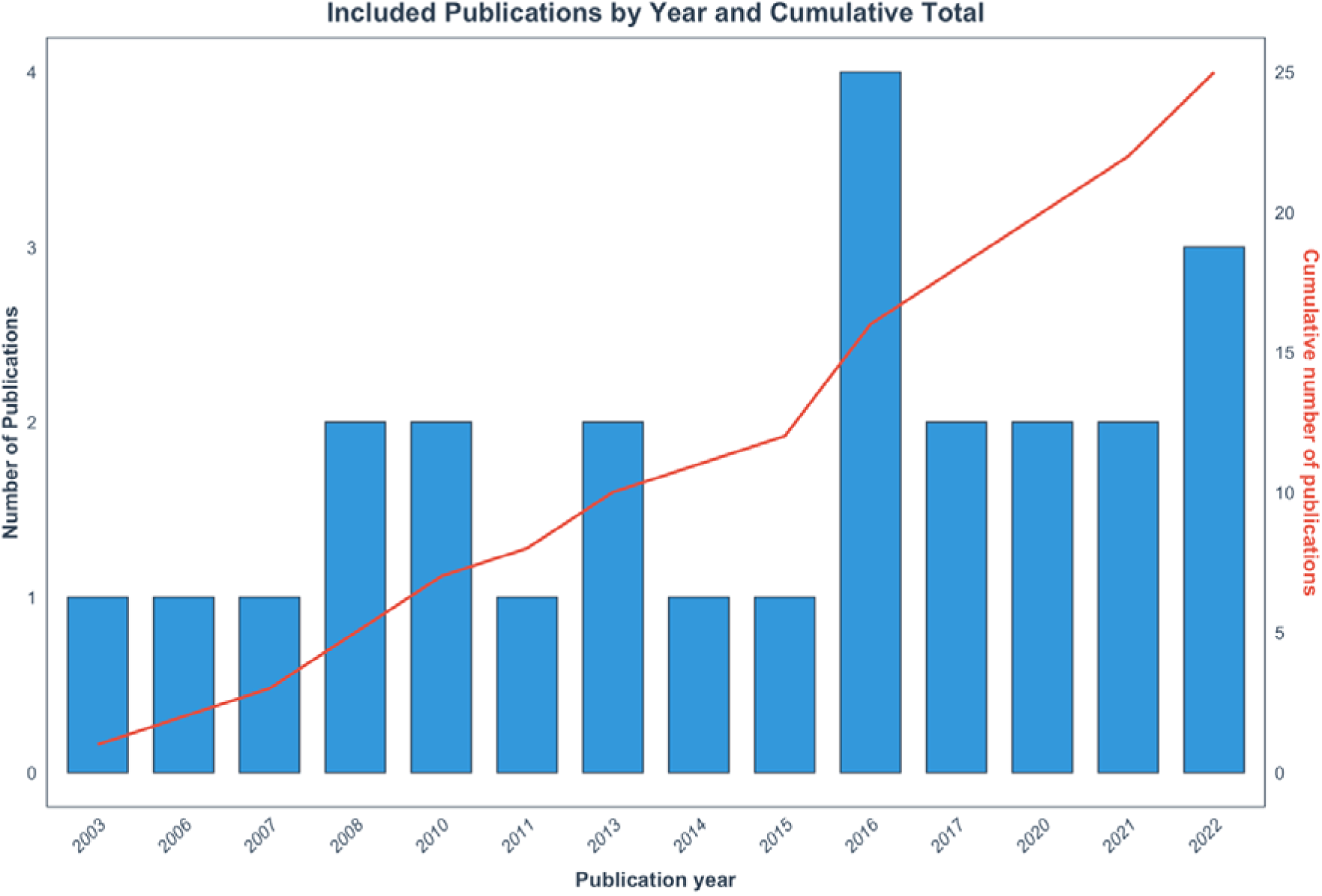
The number of publications from 2003 to 2022, with the cumulative total shown alongside. The bar chart displays the count of publications each year, while the line indicates the cumulative number of publications, scaled to fit the same axis.

### Quality of included studies

Of the 25 studies included in the review, four were classified as having a high overall risk of bias, as illustrated in Figure 3a. These studies were primarily compromised by deficiencies in the selection process and methodological rigour. Key sources of bias included small sample sizes (n < 50) and inadequately described or retrospective recruitment processes, as detailed in Supplementary Table 5. Additionally, three of these high-risk studies involved patients with lesions located in various areas, with two studies failing to account for the lesion’s location in their analyses.

**Figure 3.**
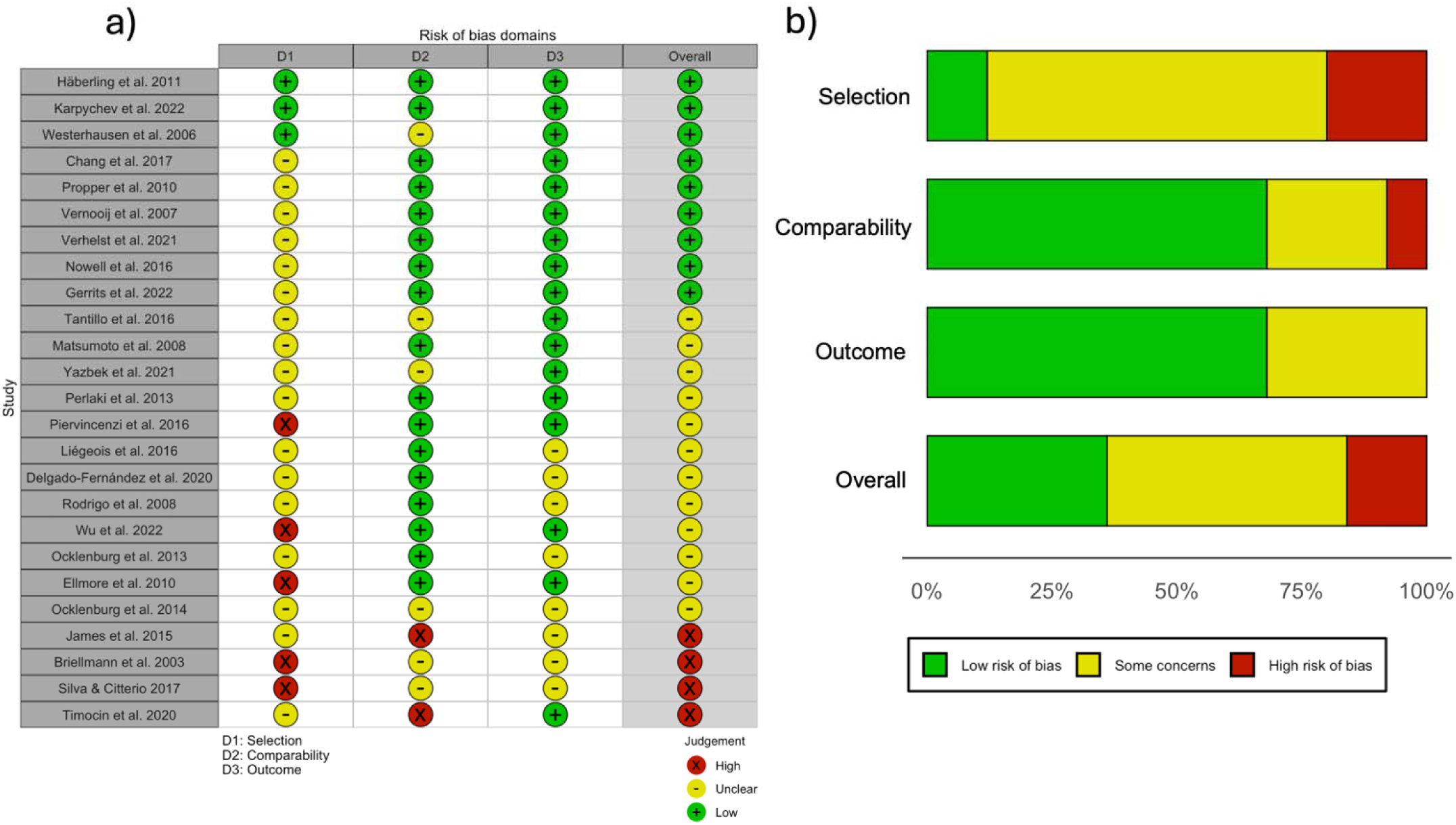
Representative table for risk of bias assessment results. a) Cochrane risk of bias assessment according to modified RoB 2 tool (Sterne et al., 2019) using traffic light plots to showcase the overall judgements for each study b) Summary plot of the overall risk of bias for each domain. The three domains of bias, including overall bias, are presented.

Methodological issues were also evident, particularly in determining functional language dominance. Problems included insufficient detail in the selection of ROIs for calculating the laterality index in functional MRI studies and a lack of justification or detail regarding the computation of this index.

Twelve studies were categorised as moderate quality, reflecting concerns similar to those found in the high-risk studies, such as cohort representation, sample size, and lesion location within clinical populations. In contrast, nine studies were rated as good quality, with only two achieving the maximum score of 11 points, as shown in Figure 3a.

Overall, the participant selection process was a significant area of concern, with only three studies attaining the highest score in this domain, as depicted in Figure 3b. Other domains showed generally strong performance, with the comparability domain having 19 studies rated without issues and only two studies scoring 1 or 0. In the outcome domain, 17 studies received full points, with no studies exhibiting high concerns and only eight studies showing some concerns.

### Participants’ information

Of the studies reviewed, 48% were conducted with diseased populations, specifically involving 12 studies with patients suffering from various conditions: eight focused on epilepsy, three on tumours, and one study included both healthy individuals and those with minor brain abnormalities. In contrast, 52% of the studies involved healthy participants. The native languages of participants varied widely, including English, Italian, Dutch, Norwegian, Russian, French, German, and Arabic. Notably, more than half of the studies (52%) did not specify the participants’ native languages. For more details, see Supplementary Table 4.

### Comparisons

The included studies employed a range of analytical methods, including correlation analyses (both Pearson’s and Spearman’s), linear regression, t-tests, non-parametric Wilcoxon tests, ANOVA, Kruskal-Wallis tests, prediction modelling, and Bayesian statistics. Among all studies, the following comparisons were used: people with left language dominance (LLD) versus right language dominance (RLD) (Delgado-Fernández et al., 2020; Gerrits et al., 2021; Matsumoto et al., 2008a; Verhelst et al., 2021; D. Wu et al., 2022), LLD versus people with atypical (bilateral+RLD) language representation (Häberling et al., 2011; Nowell et al., 2016), lateralised versus non-lateralised (Karpychev et al., 2022), strongly LLD versus mildly LLD versus RLD versus people with bilateral language representation (BLR) (Westerhausen et al., 2006), LLD versus BLR versus RLD (Tantillo et al., 2016; Timocin et al., 2020).

### Determining functional language lateralisation

Three different methods were used to determine language lateralisation. Specifically, 84% (n=21) of the studies employed fMRI for this purpose. In addition, two studies utilised the WADA test, one study applied dichotic listening, and one study combined both fMRI and WADA (Figure 4a). Regarding the ROIs used in fMRI studies, a number of brain regions were analysed, including frontal, temporal, and parietal, as well as combinations of these or whole-brain activations. In whole-brain activation studies, the ROIs referred to the activated voxels across both hemispheres, which were measured to assess brain activity during language tasks. Frontal areas were examined in 53% of the studies, temporal areas in 19%, parietotemporal areas in 16%, and whole-brain activations in 6%. One study (3%) used a combination of parietal, temporal, and frontal areas as the ROI (Yazbek et al., 2021) (Figure 4b).

**Figure 4.**
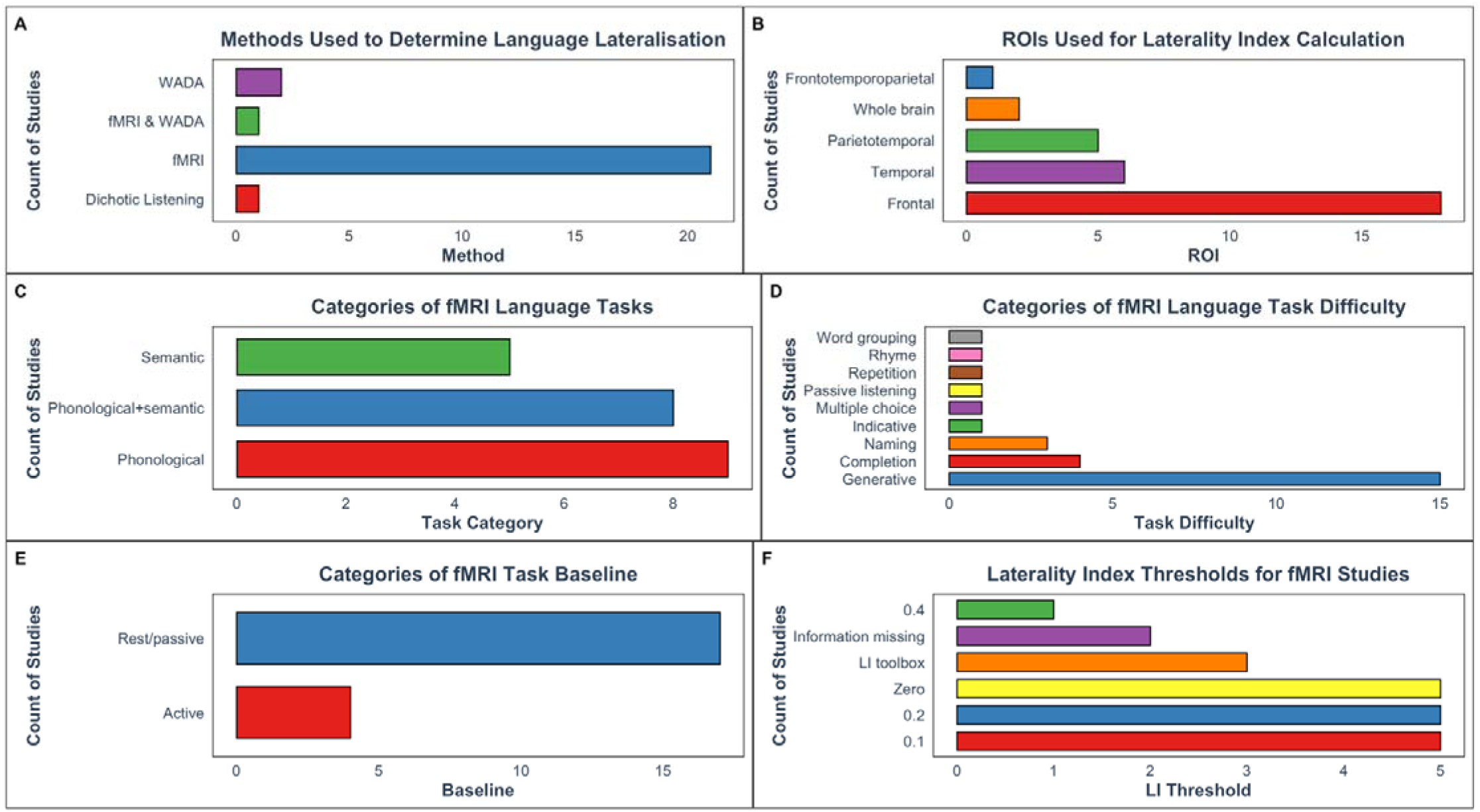
Bar charts illustrating the distribution of various aspects on language lateralisation studies. (a) Methods for determining language lateralisation include fMRI (84%, n=21), WADA (8%, n=2), dichotic listening (4%, n=1), and a combination of fMRI and WADA (4%, n=1). (b) The analysis of regions of interest (ROIs) in fMRI studies reveals that 53% of studies focused on frontal areas, 19% on temporal areas, 16% on parietotemporal areas, and 6% on whole-brain activations. Motor areas were examined in 3% of studies, and a combined parietal, temporal, and frontal ROI was used in another 3%. (c) Among fMRI tasks, 41% concentrated on phonological tasks, 36% combined phonological and semantic tasks, and 23% exclusively investigated semantic tasks. (d) For task difficulty, 54% of studies used generative tasks, 14% used word or sentence completion tasks, and 11% used naming tasks. Other tasks such as indicative questions, repetition, multiple choice, passive listening, rhyme, and word grouping were each used in a single study. (e) Baseline conditions in fMRI studies included active tasks in 19% of cases and passive conditions (fixation cross, MRI noise, or rest) in 81%. (f) Laterality index thresholds were most commonly set at 0, 0.1, and 0.2 (24% each). A laterality index toolbox was used by 14%, while 10% of studies did not report a threshold and 4% used a threshold of 0.4.

The majority of studies using fMRI to determine language lateralisation focused on phonological tasks (41%) or included both phonological and semantic tasks together (36%). A smaller proportion of studies (23%) exclusively investigated semantic tasks (Figure 4c). Regarding task difficulty, 54% of the studies employed generative tasks, where participants were required to generate words such as verbs, antonyms, words beginning with a specific letter, or semantically related words, either aloud or silently. Generative tasks are generally considered challenging. Another common method was word or sentence completion tasks, which were used in 14% of the studies reviewed. The third largest category was naming tasks, which appeared in 11% of the studies. The least common tasks included indicative (yes/no) questions, repetition, multiple choice, passive listening, rhyme tasks, and word grouping tasks, with only one study each employing these lower-difficulty tasks (Supplementary Table 4; Figure 4d). Only 4 studies (19%) used an active task as a baseline (Gerrits et al., 2022; Karpychev et al., 2022; Verhelst et al., 2021; Vernooij et al., 2007), while the remaining 81% instructed participants to either look at a fixation cross, focus on MRI noise, or rest (Figure 4e).

Lastly, we examined the thresholds used for categorising laterality groups based on the laterality index in fMRI studies. The most frequently used thresholds were 0, 0.1, and 0.2, each employed by 24% of the studies. Additionally, 14% of studies utilised a laterality index toolbox, which does not rely on a specific threshold. Two studies (10%) did not provide details on the threshold used (Propper et al., 2010; Timocin et al., 2020), and one study (4%) employed a threshold of 0.4 (Briellmann et al., 2003) (Figure 4f).

### Handedness

Fourteen studies included both right-handed and left-handed individuals (e.g. (Häberling et al., 2011; Karpychev et al., 2022; Propper et al., 2010; Vernooij et al., 2007), while one study did not provide this information (Briellmann et al., 2003). Six studies exclusively involved right-handed participants, and three studies focused solely on left-handed individuals. Given the demonstrated influence of handedness on language lateralisation (Bruckert et al., 2021; Nenert et al., 2017), we examined whether the studies accounted for handedness by either (1) including it as a covariate, or (2) analysing data separately for right- and left-handed participants. Of the studies that included both right- and left-handers, eight did not account for handedness, while seven studies did (Figure 5).

**Figure 5:**
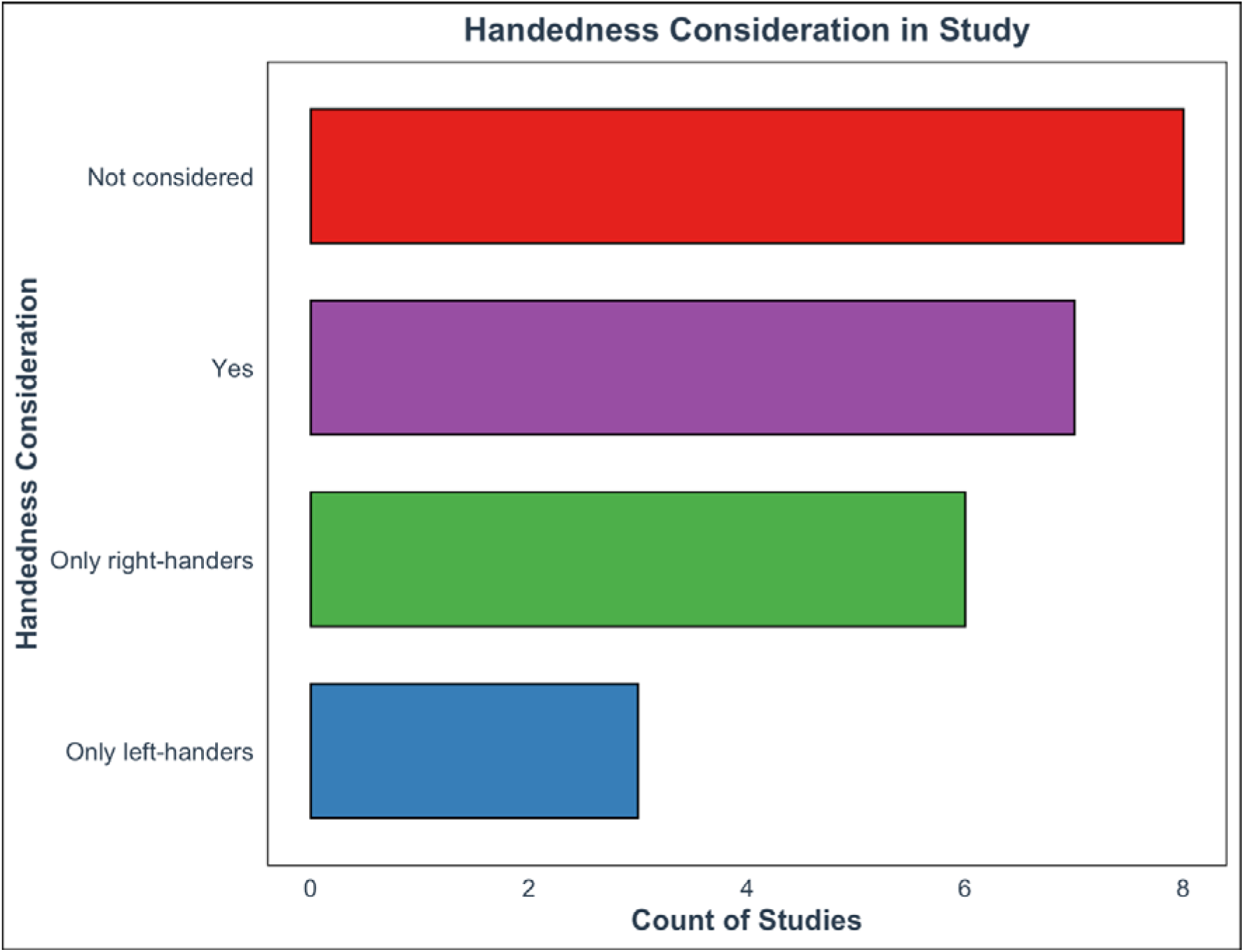
Handedness Consideration in Included Studies. This figure illustrates the frequency of studies considering handedness. The x-axis displays the categories of handedness consideration, while the y-axis represents the number of studies in each category. Bars are colour-coded according to the respective category. Categories: No: Studies that did not take handedness into consideration of laterality analyses; Yes: Studies that considered handedness either as a covariate or by separately analysing right- and left-handed participants; Only right-handers: Studies including only right-handed participants; Only left-handers: Studies including only left- handed participants.

### Diffusion MRI

Overall, two principal diffusion techniques were utilised for reconstructing white matter fibres: DTI and Constrained Spherical Deconvolution (CSD). DTI was employed in 88% of the studies, whereas only three studies (Gerrits et al., 2022; Karpychev et al., 2022; Verhelst et al., 2021) applied the more advanced CSD method (Figure 6a).

**Figure 6:**
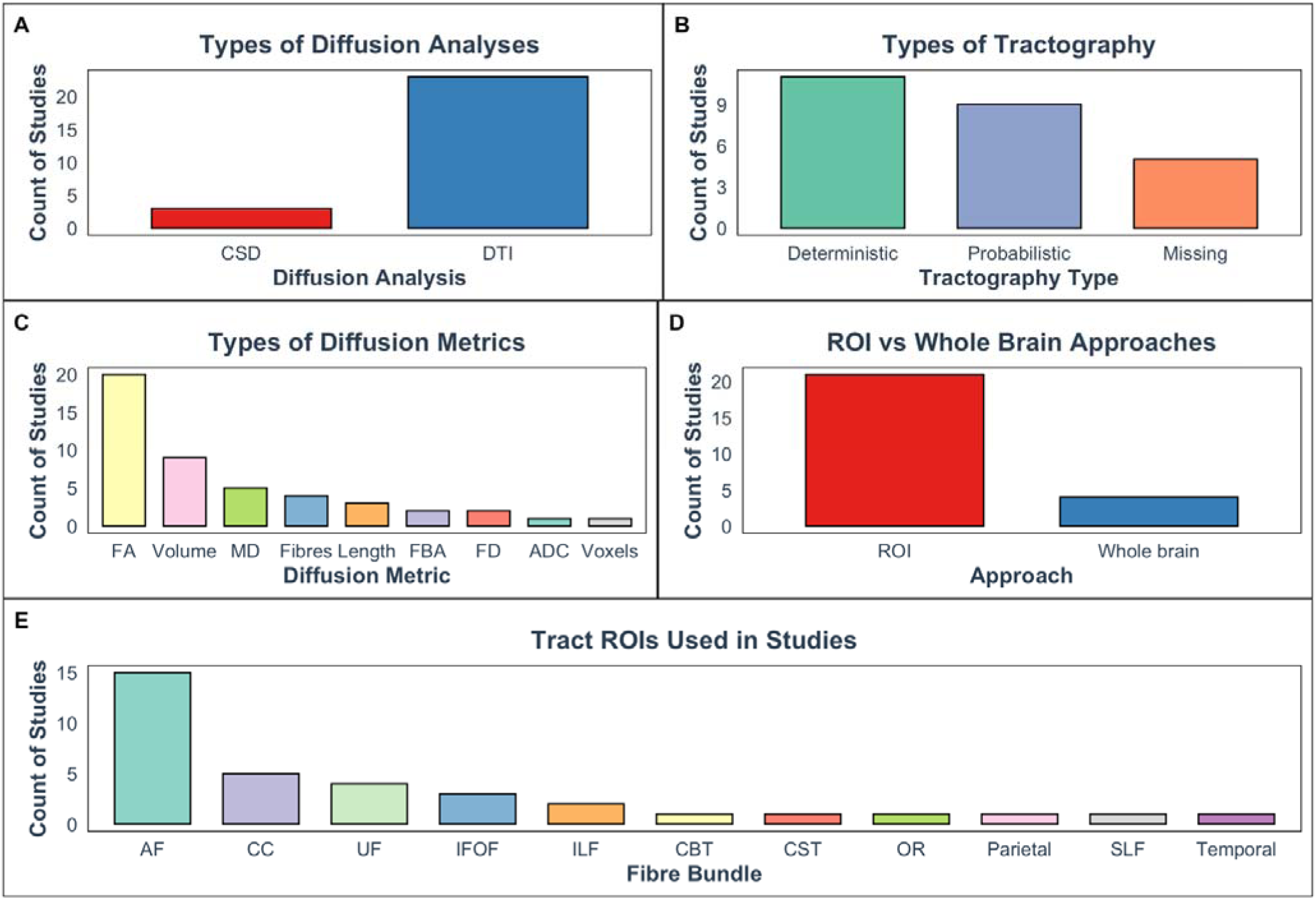
Frequency of diffusion analysis methods, tractography algorithms, diffusion metrics, ROI vs. whole-brain approaches, and brain ROIs used in the included studies. (a) Diffusion analysis methods: This panel shows the frequency of different diffusion analysis techniques used in the studies. DTI was the most commonly used method, followed by CSD. (b) Tractography algorithms: This panel illustrates the frequency of different tractography algorithms employed in the studies. Deterministic tractography was the most popular choice, followed by probabilistic tractography. A smaller proportion of studies used midsagittal seeds for tractography. (c) Diffusion metrics: This panel presents the frequency of different diffusion metrics used in the studies. FA was the most frequently used metric, followed by volume. Other metrics such as MD, number of streamlines, length, FBA, ADC, and number of voxels were less commonly used. (d) ROI vs. whole-brain approaches: This panel shows the frequency of studies that used a ROI approach compared to those that used a whole-brain approach. Most studies employed a ROI approach, selecting specific tracts in advance. (e) Brain ROIs: This panel illustrates the frequency of different brain ROIs examined in the studies. The AF was the most frequently investigated ROI, followed by the CC, UF, IFOF, and ILF. Other ROIs such as OR, SLF, CST, CBT and seeds in temporal lobe were examined in a smaller number of studies. Abbreviations: ADC: Apparent Diffusion Coefficient; AF: Arcuate Fasciculus; CBT: Corticobulbar Tract; CC: Corpus Callosum; CST: Corticospinal Tract; CSD: Constrained Spherical Deconvolution; DTI: Diffusion Tensor Imaging; FA: Fractional Anisotropy; FBA: Fixel-Based Analysis; Fibres: number of white matter streamlines; IFOF: Inferior Fronto-Occipital Fasciculus; ILF: Inferior Longitudinal Fasciculus; MD: Mean Diffusivity; Midsagittal: Midsagittal seed for tractography; Missing: tractography type undefined by the study; OR: Optic Radiation; Parietal: parietal lobe; ROI: Region of Interest; SLF: Superior Longitudinal Fasciculus; Temporal: temporal lobe; UF: Uncinate Fasciculus; Voxels: number of voxels

In terms of tractography algorithms, 44% of studies utilised a deterministic approach 36% used probabilistic tractography, and the rest five studies (20%) did not specify their tractography approach (Briellmann et al., 2003; Ocklenburg et al., 2013; Tantillo et al., 2016; Timocin et al., 2020; Westerhausen et al., 2006) (Figure 6b).

FA was the most frequently reported diffusion metric appearing in 80% of the studies. This was followed by volume of a tract, which was used in 36% of studies. Less commonly employed metrics included MD (16%), number of streamlines (12%), length and fixel-based analysis (FBA) metrics (each 8%), and apparent diffusion coefficient (ADC) and number of voxels being the least utilised (one study (4%)) (Timocin et al., 2020) (Figure 6c).

Three studies adopted a whole-brain approach (Ocklenburg et al., 2013; Perlaki et al., 2013; Verhelst et al., 2021), while the majority (88%) employed an ROI approach, with a priori selection of specific tracts (Figure 6d). Among those studies using the ROI approach, the arcuate fasciculus was the most frequently examined (68%), followed by the corpus callosum (23%), uncinate fasciculus (18%), inferior-fronto-occipital fasciculus (14%), and inferior longitudinal fasciculus (9%). Only one study investigated the corticospinal tract (Vernooij et al., 2007) (5%), optic radiation (Nowell et al., 2016) (5%), superior longitudinal fasciculus (Silva & Citterio, 2017) (5%), corticobulbar tracts (Liégeois et al., 2016) (5%), temporal ROIs (5%) (Briellmann et al., 2003), and parietal ROIs (5%) (Briellmann et al., 2003) (Figure 6e). The temporal and parietal ROIs in Briellmann et al. (2003) referred to regions within the temporal and parietal lobes used for diffusion analysis, but the study did not specify the white matter tracts involved.

Since these regions contain numerous tracts, the lack of detail limited the ability to compare it with other studies focusing on specific tracts, leading to its exclusion from further analysis.

### Arcuate fasciculus

#### Healthy population

None of the high-quality studies found associations between language laterality and asymmetry of arcuate fasciculus. However, when studies with moderate risk were considered, a few associations emerged (Figure 7). Regarding larger-scale structural properties, such as volume, length, and streamlines, there was generally no association between language lateralisation and diffusion metrics in fMRI studies (Piervincenzi et al., 2016; Propper et al., 2010). Notably, the only association was found in a dichotic listening study involving only right-handed individuals, where a significant correlation between the volume of the arcuate fasciculus and the laterality index was identified, with a larger left arcuate fasciculus associated with LLD (Ocklenburg et al., 2014). Interestingly, while not identifying a general correlation, Propper et al. (2010) found an association between the volume of arcuate fasciculus and BOLD activations in left-handed individuals, specifically in Wernicke’s area. However, it is important to note that the sample consisted solely of participants with LLD. In these individuals, a lower laterality index in Wernicke’s area indicating less left-sided lateralisation was linked to a larger right arcuate fasciculus. Similarly, Vernooij and colleagues (2007) demonstrated positive correlation between language LI in parietotemporal regions and the asymmetry of arcuate fasciculus relative fibre density (RFD) in left-lateralised right-handed individuals.

**Figure 7:**
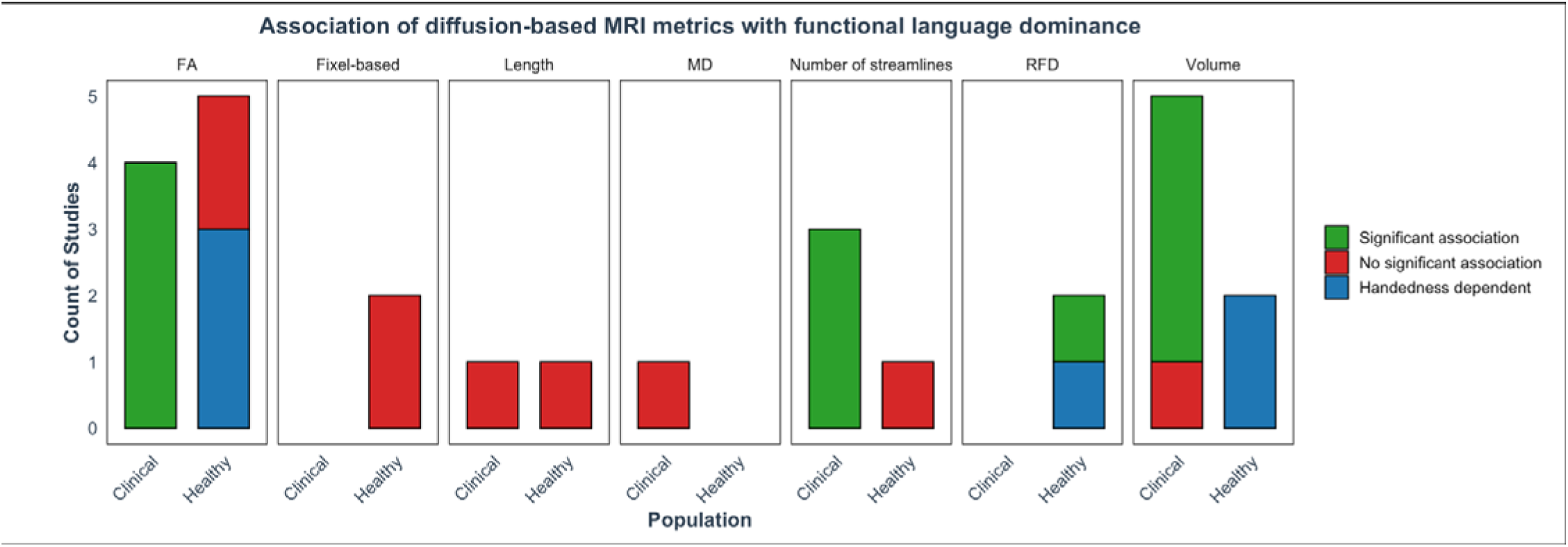
Diffusion metrics and their association with language lateralisation for the arcuate fasciculus. This plot shows the count of studies investigating various diffusion metrics (volume, length, RFD, fixel-based metrics, FA, number of streamlines) and their association with language lateralisation in healthy and clinical populations. The metrics are categorised by the presence of significant associations (“Significant association”), no significant associations (“No significant association”), or associations only evident when handedness is taken into consideration (“Handedness dependent”). For healthy populations, there is limited consistent evidence of associations, with a few notable cases related to handedness and specific study designs. In clinical populations, studies generally show a significant relationship between diffusion metrics and language lateralisation, particularly for FA and number of streamlines, highlighting the utility of these metrics in understanding language dominance.

When looking at white matter microstructure, such as FA, no association between laterality index and FA asymmetry in the arcuate fasciculus was observed in studies that did not account for handedness as a covariate (Chang et al., 2017). One study focused solely on left-handed individuals and similarly found no association (Yazbek et al., 2021). For right-handed individuals, some studies reported associations (Ocklenburg et al., 2013, 2014) while others did not (Piervincenzi et al., 2016; Yazbek et al., 2021). Among the studies that identified associations, a dichotic listening study demonstrated a significant correlation between the FA of the left arcuate fasciculus and the laterality index derived from the dichotic listening, with higher FA values linked to stronger left language dominance (Ocklenburg et al., 2014). One study found that FA in the left arcuate fasciculus positively correlated with laterality index during phonological word generation tasks involving the frontal and temporal lobes. Interestingly, passive listening to words was associated with the opposite pattern in temporal regions (i.e., higher FA in the right arcuate fasciculus correlated with greater activation in the left temporal lobe during passive listening (Ocklenburg et al., 2013)). Finally, more advanced tractography approaches, such as fixel-based analyses, did not find any associations between fixel-based metrics and language lateralisation (Gerrits et al., 2022; Verhelst et al., 2021).

These findings show no consistent association between arcuate fasciculus diffusion metrics and language laterality in healthy individuals, except in specific cases related to handedness (Figure 7a). Asymmetries were found in studies that included left lateralised people only (Propper et al., 2010; Vernooij et al., 2007), when frontal and temporal activations were examined separately (Ocklenburg et al., 2013; Vernooij et al., 2007) or in dichotic listening studies (Ocklenburg et al., 2014).

#### Clinical populations

Studies with low to moderate risk of bias found that white matter asymmetries of arcuate fasciculus in patients were linked to language lateralisation. Six studies examined clinical populations, investigating various macrostructural diffusion metrics, such as the volume, length, and number of streamlines of the arcuate fasciculus. While the number of streamlines was associated with functional language dominance (Delgado-Fernández et al., 2020; Matsumoto et al., 2008a; D. Wu et al., 2022), tract length showed no association with language dominance measured by the WADA test (Delgado-Fernández et al., 2020). Findings on volume were mixed. WADA studies identified a significant association between arcuate fasciculus volume and language dominance (Delgado-Fernández et al., 2020;

Matsumoto et al., 2008a; D. Wu et al., 2022), but one fMRI study found no association (Nowell et al., 2016). This suggests WADA may be a more reliable method to determine language dominance, making it easier to detect these relationships, possibly due to the arbitrary choice of ROIs for LI computation. For example, Nowell et al. (2016) only looked at frontal activations and did not consider temporal regions. When studies with a higher risk of bias were considered, an fMRI study (Silva & Citterio, 2017) investigating both phonological and semantic aspects of language found a significant association between structural asymmetries and language dominance, regardless of lesion location. This suggests that the volume of the arcuate fasciculus may be related to language lateralisation in clinical populations, independent of factors such as handedness and lesion size.

All studies investigating the relationship between arcuate fasciculus FA and language lateralisation found significant associations. Epilepsy studies using WADA to determine language dominance (Figure 7) (Delgado-Fernández et al., 2020; Ellmore et al., 2010; D. Wu et al., 2022) showed that the language-dominant hemisphere had significantly higher FA in the arcuate fasciculus compared to non-dominant hemisphere. Other studies using fMRI focused on different epilepsy types. One study examined left and right temporal lobe epilepsy separately (Rodrigo et al., 2008), while another focused solely on patients with left temporal lobe epilepsy (Chang et al., 2017). Interestingly, Chang et al. (2017) found greater arcuate fasciculus FA asymmetry in right-lateralised individuals, with higher FA in the right arcuate fasciculus in the temporo-parietal and inferior frontal regions among those with left temporal lobe epilepsy. Similarly, Rodrigo et al. (2008) showed positive relationship between arcuate fasciculus FA and LI in Broca’s area in patients with right temporal lobe epilepsy. Finally, MD was not associated with language lateralisation (Rodrigo et al., 2008).

#### Corpus callosum

Studies on the corpus callosum were approximately three times less frequent than those on the arcuate fasciculus (Figure 6e). Among all studies examining the corpus callosum, five focused on the healthy populations (Häberling et al., 2011; Karpychev et al., 2022; Perlaki et al., 2013; Verhelst et al., 2021; Westerhausen et al., 2006) while only two examined individuals with brain tumours (Tantillo et al., 2016; Timocin et al., 2020). Most studies on healthy individuals were of high quality, with only one study classified as having a moderate risk of bias (Perlaki et al., 2013). However, studies on clinical populations tended to have a higher risk of bias, with moderate to severe risk identified in both tumour studies (Tantillo et al., 2016; Timocin et al., 2020).

#### Healthy population

In healthy populations, findings varied across diffusion metrics and regions. For example, studies that examined the corpus callosum as a whole found that higher FA was associated with BLR or RLD (Häberling et al., 2011). Regarding specific regions, studies that segmented the corpus callosum showed different results. For example, Westerhausen et al. (2006) combined the genu (anterior part) and splenium (posterior part) and observed that strongly left-lateralised individuals exhibited lower MD, but this was only evident in those with strong left-lateralisation. Posterior sections of the corpus callosum showed mixed results across studies. While some studies found no associations for FA (Häberling et al., 2011; Karpychev et al., 2022) and volume (Westerhausen et al., 2006), others reported that greater volume (Karpychev et al., 2022) and higher relative anisotropy (Westerhausen et al., 2006) in the posterior corpus callosum correlated with increased left-lateralisation.

In whole-brain studies, conventional DTI was insufficient for detecting corpus callosum associations with language lateralisation (Perlaki et al., 2013). More advanced tractography techniques, such as fixel-based analysis, showed a significant association between the right forceps minor, which is an anterior commissural tract, and language lateralisation (Verhelst et al., 2021). Authors found that a larger fibre cross-section (FC) in the right forceps minor was associated with right language dominance, while the opposite pattern was seen in left-dominant individuals. However, this study did not include people with BLR.

#### Clinical population

In clinical populations, larger corpus callosum volumes was associated with atypical language lateralisation were generally observed. Tantillo et al. (2016) found that greater FA in the anterior corpus callosum was associated with BLR in Broca’s area. Similarly, more recent studies showed that higher number of tracts in corpus callosum was associated with more atypical (right and bilateral) representation of language (Timocin et al., 2020). However, Timocin et al. (2020) did not divide the corpus callosum to different parts or provide lateralisation in temporal and frontal lobes separately, limiting comparisons.

#### Other association tracts

Four other white matter bundles were described in the studies: the superior longitudinal fasciculus, inferior longitudinal fasciculus, uncinate fasciculus, and inferior fronto-occipital fasciculus. The left superior longitudinal fasciculus was associated with typical left language lateralisation in healthy individuals, as shown by higher FA in this tract (Perlaki et al., 2013). In contrast, studies on epilepsy patients found no significant relationship between the volume of the superior longitudinal fasciculus and language lateralisation (Silva & Citterio, 2017), though this finding should be interpreted with caution due to methodological concerns. Similarly, the inferior longitudinal fasciculus exhibited higher relative fibre density in the left hemisphere in healthy individuals, correlating with left lateralisation (James et al., 2015), while clinical studies did not observe this association (Ellmore et al., 2010).

Regarding the right uncinate fasciculus, both volume and FA were associated with left language dominance in whole-brain and dichotic listening studies in healthy populations (Ocklenburg et al., 2013; 2014). In contrast, relative fibre density showed no relationship with language dominance (James et al., 2015). Clinical studies using the WADA test to determine language dominance found no associations between FA and language laterality (Delgado-Fernández et al., 2020; Ellmore et al., 2010), nor between the number of fibres and language laterality (Delgado-Fernández et al., 2020). The only measures to show positive associations with left uncinate fasciculus and left language dominance were fibre length and volume, although this was reported in only one study (Delgado-Fernández et al., 2020).

The inferior fronto-occipital fasciculus was examined in two studies on healthy individuals and two in people with epilepsy, yet none of these studies identified significant associations between diffusion metrics including FA, MD, or relative fibre density and language lateralisation measured in phonological or semantic tasks (Chang et al., 2017; James et al., 2015; Rodrigo et al., 2008)

Overall, FA was a strong predictor of language lateralisation in the left superior longitudinal fasciculus and uncinate fasciculus in healthy individuals. For clinical populations, however, only the volume of the uncinate fasciculus consistently showed a significant link to language lateralisation. Larger volumes of this tract were associated with increased left language dominance in both healthy and clinical groups. On the other hand, other tracts and diffusion metrics in clinical populations did not show consistent patterns with language dominance.

Projection tracts

Several studies have examined projection tracts, including the corticospinal tract, corticobulbar tract, and optic radiation. Studies report no significant association between language lateralisation and the corticospinal or corticobulbar tracts in healthy individuals (Liégeois et al., 2016; Vernooij et al., 2007). In contrast, Nowell et al. (2016) studied optic radiation, specifically Meyer’s loop, in people with epilepsy. Their findings demonstrated that the spatial position of Meyer’s loop relative to the temporal lobe was associated with language dominance, with a more anterior position in left-lateralised individuals and a more posterior position in right-lateralised individuals.

## Discussion

In this review, we investigated the relationship between functional language dominance and diffusion MRI-based white matter measures, considering factors such as methods for determining language lateralisation (e.g., WADA, fMRI, dichotic listening), ROIs used for calculating laterality indices, the nature of language tasks (e.g., phonological or semantic focus, task difficulty, and baseline conditions), laterality index thresholds, diffusion analysis techniques (DTI or CSD), types of tractography, diffusion metrics, diffusion MRI statistical approaches (ROI-based or whole-brain), and modulating factors such as handedness and lesion location. Our findings revealed that (a) arcuate fasciculus FA showed a strong association with language lateralisation in clinical populations but not in healthy individuals, where results were more heterogeneous and often influenced by handedness; (b) other white matter tracts, including the corpus callosum, uncinate fasciculus, inferior fronto-occipital fasciculus, and inferior longitudinal fasciculus, did not show consistent associations with language lateralisation; (c) semantic tasks that utilised temporal ROIs in fMRI studies were more strongly associated with diffusion MRI measures compared to other task types; and (d) major concerns in the studies included variability in sample selection, heterogeneity in language fMRI and diffusion protocols, and inconsistencies in analytical approaches, all of which made it difficult to compare findings across studies and draw robust conclusions, highlighting the urgent need for standardised methodologies.

### Arcuate fasciculus

The relationship between language lateralisation and diffusion metrics of the arcuate fasciculus is not consistently demonstrated, particularly when comparing clinical and healthy populations. In clinical populations, particularly individuals with epilepsy, higher FA within the arcuate fasciculus is strongly associated with LLD. However, this relationship is less consistently observed in healthy populations. Chang et al. (2017) argue that this variability could be due to the small number of atypically lateralised individuals in healthy cohorts. This stands in stark contrast to clinical populations, which often exhibit a much broader range of laterality indices (Tailby et al., 2017).

Studies focused on healthy controls may therefore lack the statistical power needed to produce robust findings on patterns of lateralisation. Nonetheless, even studies with a substantial proportion of left-handed participants—who are more likely to exhibit atypical language dominance—have failed to find a significant relationship between diffusion-based MRI measures of the arcuate fasciculus and language lateralisation (e.g., Gerrits et al., 2022; Verhelst et al., 2021; Vernooij et al., 2007). This discrepancy raises questions about the underlying factors driving these inconsistencies, suggesting that the mere presence of atypical individuals may not fully explain the lack of consistent findings.

It is important to differentiate between atypical lateralisation due to brain injury and that which occurs naturally in healthy individuals. In clinical populations, such as those with epilepsy or stroke, atypical lateralisation often results from neuroplastic adaptations. For example, language dominance may shift to the right hemisphere in epilepsy, leading to increased FA in the right arcuate fasciculus, while in stroke, increased FA in the left arcuate fasciculus reflects a compensatory response to left hemisphere damage (Chang et al., 2017; Forkel et al., 2014). These findings suggest that neuroplasticity, driven by the brain’s need to compensate for functional deficits, leads to structural adaptations in white matter. In contrast, the neural system in healthy individuals operates more efficiently, thereby reducing the necessity for compensatory increases in FA. This aligns with the notion that individuals with higher cognitive capacity exhibit less neural activation during task performance (Prat, 2011). For example, high-vocabulary readers show reduced activation in language regions compared to low- vocabulary readers during fMRI tasks (Prat, 2011). Similarly, tasks that demand greater semantic processing often result in activation not only in the dominant hemisphere but also in the contralateral hemisphere (Cosgrove et al., 2023; Jung et al., 2021; Wu & Hoffman, 2023). This flexibility mirrors the brain’s ability to compensate in the face of perturbation or brain damage, highlighting the resilience-related mechanisms of semantic cognition. While dynamic shifts in neural activity have been observed in both healthy individuals and those with brain injury in fMRI studies, there has been little research exploring the relationship between these shifts, hemispheric language asymmetry, and white matter organisation. More work is needed to investigate how task difficulty influences these adaptive changes in language lateralisation.

It is also possible that healthy individuals with atypical language dominance activate alternative neural networks (in contrast to those observed in clinical populations), resulting in more complex activation patterns during fMRI tasks. People with BLR, for instance, often recruit additional areas such as the angular gyrus and middle temporal gyrus, which are less commonly activated in left-lateralised individuals (Bidula et al., 2017). Conversely, people with RLD tend to exhibit mirror-image activations in the left hemisphere (Bidula et al., 2017). These distinct activation patterns may complicate the relationship between FA and lateralisation, particularly when studies assume homogeneity across different lateralisation types.

Focusing exclusively on individuals with LLD reveals a more consistent association: higher FA in the left arcuate fasciculus is linked to stronger left hemisphere dominance, while lower FA suggests weaker dominance (Propper et al., 2010; Vernooij et al., 2007). However, in clinical populations, such as those with epilepsy, associations between white matter structure and lateralisation appear more robust across different lateralisation types, including BLR and RLD. The reason for this difference is still unclear, though it is possible that clinical populations have less complex language organisation (i.e. utilising mirror regions regardless of lateralisation), a hypothesis that remains unexplored.

### Commissural tracts

Two primary models describe the role of the corpus callosum in language lateralisation: the inhibitory and excitatory models (Bloom & Hynd, 2005). The inhibitory model suggests that in lateralised individuals, the corpus callosum facilitates hemispheric connectivity by suppressing activity in the non-dominant hemisphere, thus enhancing lateralisation. In contrast, the excitatory model posits that, in bilaterally dominant individuals, the corpus callosum supports information transfer between both hemispheres (Westerhausen et al., 2006). Our findings offer mixed evidence regarding the role of the corpus callosum in language lateralisation. Similar to the arcuate fasciculus, diffusion metrics within the corpus callosum show a more pronounced association with language lateralisation in clinical populations. In these groups, larger corpus callosum size and greater atypicality in language lateralisation are consistently observed (Tantillo et al, 2016; Timocin et al., 2020). This suggests adaptive plasticity, where increased interhemispheric communication strengthens white matter connectivity to support compensatory processes for maintaining or recovering language function (Hebb, 2002). In healthy individuals, however, these relationships appear less distinct, with the corpus callosum potentially operating under a more dynamic balance of inhibitory and excitatory functions depending on task demands and environmental factors (Ocklenburg & Guo, 2024).

This highlights a key distinction: in clinical populations, the corpus callosum may prioritise compensatory excitatory processes to support language functions, whereas in healthy populations, it may rely on flexible mechanisms aimed at maintaining interhemispheric balance and optimising efficiency.

### ROI selection

The selection of ROIs in functional studies can significantly impact the findings from language lateralisation studies. For instance, focusing on temporal regions for semantic tasks has consistently yielded robust results for determining language lateralisation (Chang et al., 2017; Karpychev et al., 2022; Propper et al., 2010; Vernooij et al., 2007). Instead, studies focusing on frontal regions’ activity during phonological tasks have produced mixed findings (Ellmore et al., 2010; Gerrits et al., 2021). One potential explanation for the variability in Broca’s area may stem from the lack of consensus regarding its anatomical boundaries, which can lead to misinterpretations of its role in language lateralisation (Tremblay & Dick, 2016). Additionally, cases in which resection of Broca’s area did not result in postoperative language deficits challenge traditional models of its role in language processing, suggesting a more flexible functional architecture (Benzagmout et al., 2007; Duffau et al., 2008; Lubrano et al., 2010; Sanai et al., 2008). This flexibility raises important questions about the capacity of conventional models of language lateralisation to account for individual variability in brain anatomy and function.

Current gaps and future directions

This systematic review highlights several limitations that could guide future research in this field. One key issue is that language lateralisation is not a static characteristic; rather, it fluctuates depending on the nature of the task. As Parker et al. (2022) highlight, lateralisation can vary significantly with task demands. Studies using both speech perception and generation tasks have demonstrated that lateralisation often depends on the task, particularly for individuals with atypical lateralisation (Woodhead et al., 2021). This underscores the importance of using diverse tasks when assessing language lateralisation, especially in clinical contexts where accurate mapping is critical for presurgical planning.

The segmentation of tracts is another critical factor. The anterior and posterior segments of the arcuate fasciculus have been linked to distinct language functions (Catani & Dawson, 2017), although only one study (Matsumoto et al., 2008b) looked into associations between separate segments. Failing to distinguish between these segments may obscure important functional differences, as aggregating data across the entire tract could mask associations specific to phonological or semantic processes. Dávolos et al. (2020) further demonstrate that right hemisphere tracts become more engaged during lower-demand tasks, while higher-demand tasks (such as semantic retrieval) predominantly activate left hemisphere tracts. This highlights how task difficulty and cognitive load influence hemispheric activation, complicating the study of language lateralisation.

Future research should incorporate fine-grained analyses of tract segmentation to better understand these mechanisms and avoid oversimplifying the role of major white matter tracts like the arcuate fasciculus.

Sample size also poses a significant challenge. The study with the largest sample size was 89 people (Westerhausen et al., 2006), which is problematic because, to decipher structural correlates in atypically lateralised individuals, a study needs to be more robust, requiring a substantial number of atypical cases. Small sample sizes ultimately result in a limited number of atypical cases (Westerhausen et al., 2014), which reduces the statistical power and generalisability of the findings, making it difficult to detect meaningful differences or draw reliable conclusions about atypical populations.

Methodological shortcomings further hinder progress in this field. 88% of the diffusion studies employed the ROI approach, which increases the chances of false positives and limits the generalisability of the research due to variations in how ROIs are defined (e.g., anatomical location or tract segmentation vs. whole). This also restricts the identification of ’less conventional tracts,’ with a heavy emphasis on the arcuate fasciculus (in 68% of studies), despite the growing recognition that language processing involves a wide network of interactions (Thiebaut de Schotten & Forkel, 2022). Lastly, none of the studies explored the impact of task difficulty on the structure-function relationship of language lateralisation, which might explain the absence of associations in healthy individuals.

To address these limitations, future studies should aim to recruit larger sample sizes, standardise ROI selection methods, and incorporate a variety of tasks to better understand the complexity of language lateralisation.

### Theoretical and clinical implications

Our findings may enhance the theoretical understanding of how white matter structure contributes to language lateralisation, while also highlighting significant gaps in the current literature (Bookheimer, 2007). Notably, more than two-thirds of the studies in our review focused on the arcuate fasciculus, highlighting the need for language lateralisation research to move beyond traditional Broca-Wernicke models (Tremblay & Dick, 2016). Recent studies showed that less than 5% of white matter fibres in the middle part of the arcuate fasciculus directly connected Broca’s and Wernicke’s areas (Rosen & Halgren, 2022), which suggests that information between such areas might be transmitted either directly via long-range or indirect connections within other fibre tracts (Mollon et al., 2022). Furthermore, recent functional MRI studies indicate that BOLD activations during language task can be predicted by resting-state fMRI data through connectivity analyses (Braga et al., 2020). This approach outperforms regional mask methods that focus solely on Broca’s and Wernicke’s areas, where language lateralisation is particularly challenging to predict in atypically lateralised individuals (Phillips et al., 2021; Rolinski et al., 2020). These findings highlight an urgent need to adopt more comprehensive, connectivity-based analytical methods to better capture the complexity of language processing and lateralisation, particularly in atypical populations.

Advancing the theoretical understanding of white matter structure and language dominance is essential for improving clinical research, particularly in presurgical evaluations of individuals with brain lesions. Although fMRI remains the gold standard for assessing language lateralisation (Dym et al., 2011), its slower acquisition time compared to DTI poses challenges, particularly in populations where extended immobility, cognitive impairment and chronological age is a significant issue. Should the relationship between DTI metrics and language dominance prove robust, DTI could emerge as a viable, cost-effective alternative for presurgical evaluation. Ultimately, this synthesis of literature will contribute to a more nuanced understanding of the neuroanatomical correlates of language processing and may inform future research directions in the field of cognitive neuroscience.

## Conclusions

The relationship between white matter structure and language lateralisation exhibits significant differences between clinical populations, such as individuals with epilepsy, and healthy individuals, primarily due to methodological challenges. Clinical populations demonstrate a clear association between diffusion metrics—such as FA and mean diffusivity— and language lateralisation, reflecting structural adaptations linked to atypical language dominance. In contrast, healthy individuals often show less consistent correlations, which may result from subtler structural changes, variability in neural activation patterns, and methodological issues like ROI selection and task demands. Studies indicate that ROI selection can notably influence findings, with temporal regions typically yielding more robust results than frontal regions. Furthermore, the segmentation of white matter tracts and the specific language tasks employed are crucial for understanding these relationships.

## CRediT authorship contribution statement

Ieva Andrulyte: Conceptualisation, writing, reviewing all papers, original draft Space for other authors

