## Supplementary Table 2 for "Does white matter structure relate to hemispheric language lateralisation? A systematic review"

Broad electronic searches will include terms related to language lateralization and white matter asymmetries and will be conducted using free text words/phrases with no language or date restrictions in the following databases:

**Table 1. The search strategy used for Ovid MEDLINE investigation**

| # | Searches |
| --- | --- |
| 1. | "language lateral*".mp. |
| 2. | "language dominance".mp. |
| 3. | "language asymmetry$".mp. |
| 4. | 1 or 2 or 3 |
| 5. | language.mp. |
| 6. | "hemispheric dominance".mp. |
| 7. | "hemispheric asymmetry$".mp. |
| 8. | "brain asymmetry$".mp. |
| 9. | 6 or 7 or 8 |
| 10. | 5 and 9 |
| 11. | 4 or 10 |
| 12. | DTI.mp. |
| 13. | "diffusion tensor imaging".mp. |
| 14. | tractography.mp. |
| 15. | "white matter".mp. |
| 16. | fibre$.mp. |
| 17. | fiber$.mp. |
| 18. | 12 or 13 or 14 or 15 or 16 or 17 |
| 19. | 11 and 18 |

* *mp = title, book title, abstract, original title, name of substance word, subject heading word, floating sub-heading word, keyword heading word, organism supplementary concept word, protocol supplementary concept word, rare disease supplementary concept word, unique identifier, synonyms*

**Table 2. The search strategy used for Scopus investigation**

| # | Searches |
| --- | --- |
| 1. | ( ( TITLE-ABS-KEY ( "language lateral*" OR "language dominance" OR "language asymmetry$" ) ) OR ( TITLE-ABS-KEY ( "language" AND ( "hemispheric dominance" OR "hemispheric asymmetry$" OR "brain asymmetry$") ) ) ) AND ( TITLE-ABS-KEY ( "dti" OR "diffusion tensor imaging" OR "tractography" OR "white matter" OR "fibre$" OR "fiber$" ) ) |

** TITLE-ABS-KEY = article title, abstract, keywords*

**Table 3. The search strategy used for Web of Science investigation**

| # | Searches |
| --- | --- |
| 1.; 2.; 3.; 4.; 5.; 6.; 7.; 8.; 9.; 10.; 11.; 12.; 13.; 14.; 15.; 16.; 17.; 18.; 19. | "language lateral*” (Topic); "language dominance” (Topic); "language asymmetry$” (Topic); #1 OR #2 OR #3; language (Topic); "hemispheric dominance” (Topic); "hemispheric asymmetry$” (Topic); "brain asymmetry$” (Topic); #6 OR #7 OR #8; #5 AND #9; #4 OR #10; DTI (Topic); "diffusion tensor imaging” (Topic); tractography (Topic); "white matter” (Topic); fibre$ (Topic); fiber$ (Topic); #12 OR #13 OR #14 OR #15 OR #16 OR #17; #18 AND #11 |

**Topic = title, abstract, author keywords, and Keywords Plus.*
